# Human Sensitivity to Perturbations Constrained by a Model of the Natural Image Manifold

**DOI:** 10.1101/320531

**Authors:** Ingo Fruend, Elee Stalker

## Abstract

Humans are remarkably well tuned to the statistical properties of natural images. However, quantitative characterization of processing within the domain of natural images has been difficult because most parametric manipulations of a natural image make that image appear less natural. We used generative adversarial networks (GANs) to constrain parametric manipulations to remain within an approximation of the manifold of natural images. In the first experiment, 7 observers decided which one of two synthetic perturbed images matched a synthetic unperturbed comparison image. Observers were significantly more sensitive to perturbations that were constrained to an approximate manifold of natural images than they were to perturbations applied directly in pixel space. Trial by trial errors were consistent with the idea that these perturbations disrupt configural aspects of visual structure used in image segmentation. In a second experiment, 5 observers discriminated paths along the image manifold as recovered by the GAN. Observers were remarkably good at this task, confirming that observers were tuned to fairly detailed properties of an approximate manifold of natural images. We conclude that human tuning to natural images is more general than detecting deviations from natural appearance, and that humans have, to some extent, access to detailed interrelations between natural images.

## Introduction

The images that we experience in our everyday visual environment are highly complex and our visual system seems to be adapted to perform well with these natural images. However, they only comprise a small part of all possible digital images (Simoncelli & Olshausen, 2001; Geisler, 2008).

Humans are very quick at making simple decisions in natural images (Thorpe et al., 2001). We can easily complete missing image components with the most natural looking value (Bethge et al., 2007) or detect visual deviations from naturalness (Gerhard et al., 2013; Wallis et al., 2017; Fründ & Elder, 2013). This precise tuning seems to be mostly restricted to foveal areas and humans are much less sensitive to deviations from naturalness in the periphery (Wallis & Bex, 2012). In fact, two physically different images that match in only a coarse set of image statistics in the periphery, typically appear to be the same (Freeman & Simoncelli, 2011).

Most of these studies are concerned with human sensitivity to deviations from naturalness. It is however, less clear how to characterize human performance within the manifold of natural images. One challenge is that most experimental manipulations of natural images make the image itself appear less natural. For example, Bex (2010) manipulates images by local deformations. He finds that sensitivity to these deformations is tuned to the spatial frequency at which they occur. Although the deformations used by Bex (2010) only moderately alter the power spectrum of natural images, they considerably disrupt higher order statistical properties such as the phase spectrum of the images (Wichmann et al., 2006). Others, have manipulated images by introducing “dead leaves”—small homogeneous patches—in different locations (Wallis & Bex, 2012) or manipulating the correlation structure of the images (McDonald & Tadmor, 2006). For small patches presented in the visual periphery, observers can often not detect these manipulations. However, we argue that these manipulations still make the image appear less natural rather than studying visual processing within the domain of natural images. A possible solution to this problem is to use selective sampling (Sebastian et al., 2017): Instead of attempting to manipulate the image directly, one chooses another natural image that *by chance* shows the desired manipulation. Although this approach guarantees that the resulting “manipulations” remain natural, it is highly dependent on the indexing mechanism used to select the “manipulated image”. The selective sampling approach is dependent on the indexing mechanism used because for any new feature, a new indexing mechanism would need to be implemented. More importantly, if there aren’t sufficiently many exemplars for a given feature, a new database would be needed. This dependence creates a limitation on generalizing the selective sampling approach to higher levels of visual processing.

Recent advances in machine learning might provide a means to constrain image manipulations to the domain of natural images. Here, we focus on a class of very powerful generative image models, known as generative adversarial nets (GANs, Goodfellow et al., 2014; Radford et al., 2016; Arjovsky et al., 2017; Gulrajani et al., 2017; Miyato et al., 2018; Hjelm et al., 2018). GANs learn a mapping—called the generator—from an isotropic Gaussian distribution to the space of images. One defining feature of GANs is the use of an auxiliary classification function—often called the critic—to judge how good the generator mapping is. Specifically, the critic attempts to predict if a given image has been generated by mapping isotropic Gaussian noise through the generator, or if the image is an instance from the training database. Generator and critic are trained in alternation, where the generator is trained to increase the errors of the critic and the critic is trained to decrease its own error (see for example Goodfellow et al., 2014, for details). In general, generator and critic can be any possible transformation, but they are typically implemented as artificial neural networks with multiple hidden layers (Goodfellow et al., 2014; Radford et al., 2016). Although never studied quantitatively, images generated from GANs look quite similar to natural images and manipulations in a GAN’s latent space seem to correspond in a meaningful way to perceptual experience. For example, Radford et al. (2016) start with a picture of a smiling woman, subtract the average latent representation of a neutral woman’s face and add a neutral man’s face to arrive at a picture of a smiling man. Similarly, Zhu et al. (2016) illustrate that projecting perceptually meaningful constraints back to a GAN’s latent space allows creation of random images with specified features (e.g. edges or colored patches) in the specified locations. Together, this suggests that GANs recover a reasonably good approximation to the manifold of natural images.

In this study, we used GANs to manipulate generated images and observers made perceptual judgments about these images. In experiment 1, the observers viewed a target image and decided which one of two noisy comparison images corresponded to that target. Crucially, noise was either applied directly in pixel space or it was restricted to remain within an approximation to the manifold of natural images by applying it in the latent space of a GAN. We found that this task was more difficult if noise was applied in the latent space of a GAN, suggesting that noise in the GAN’s latent space actually changes image features that are relevant for image recognition, while noise that was directly applied in pixel space only resulted in degradation of the image without necessarily changing image content. In experiment 2, observers were asked to detect changes of direction in videos that were constructed by moving along smooth paths through a GAN’s latent space and we found that observers performed significantly above chance even for very small directional changes, suggesting that GANs not only recover a good approximation to the manifold of natural images, but that they also recover a perceptually meaningful parameterization of this manifold.

## Experiment 1: Sensitivity to perturbations within the approximate natural image

### Method

#### Training generative adversarial nets

We trained a Wasserstein-GAN (Arjovsky et al., 2017) on the 60 000 32 × 32 images contained in the CIFAR10 dataset (Krizhevsky, 2009) using gradient penalty as proposed by (Gulrajani et al., 2017). See Figure 1A for example training images. In short, a GAN consists of a generator network *G* that maps a latent vector *z* to image space and a critic network *D* that takes an image as input and predicts whether that image is a real image from the training dataset or an image that was generated by mapping a latent vector through the generator network (see Figure 2 and Gulrajani et al., 2017 for details of the architecture of the two networks). The generator network and the critic network were trained in alternation using stochastic gradient descent. Specifically, training alternated between 5 updates of the critic network and one update of the generator network. Updates of the critic network were chosen to minimize the loss

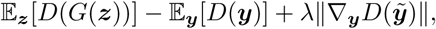
and updates of the generator were chosen to maximize this loss. Here, 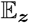 and 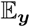 denote averages over a batch of 64 latent vectors ***z*** or training images ***y*** respectively. Furthermore, ∇***_y_*** denotes the gradient with respect to image pixels ***y***, which was evaluated at random points along straight line interpolations between real and generated images (see Gulrajani et al., 2017 for details). We set *λ* = 10 during training.

**Figure 1:**
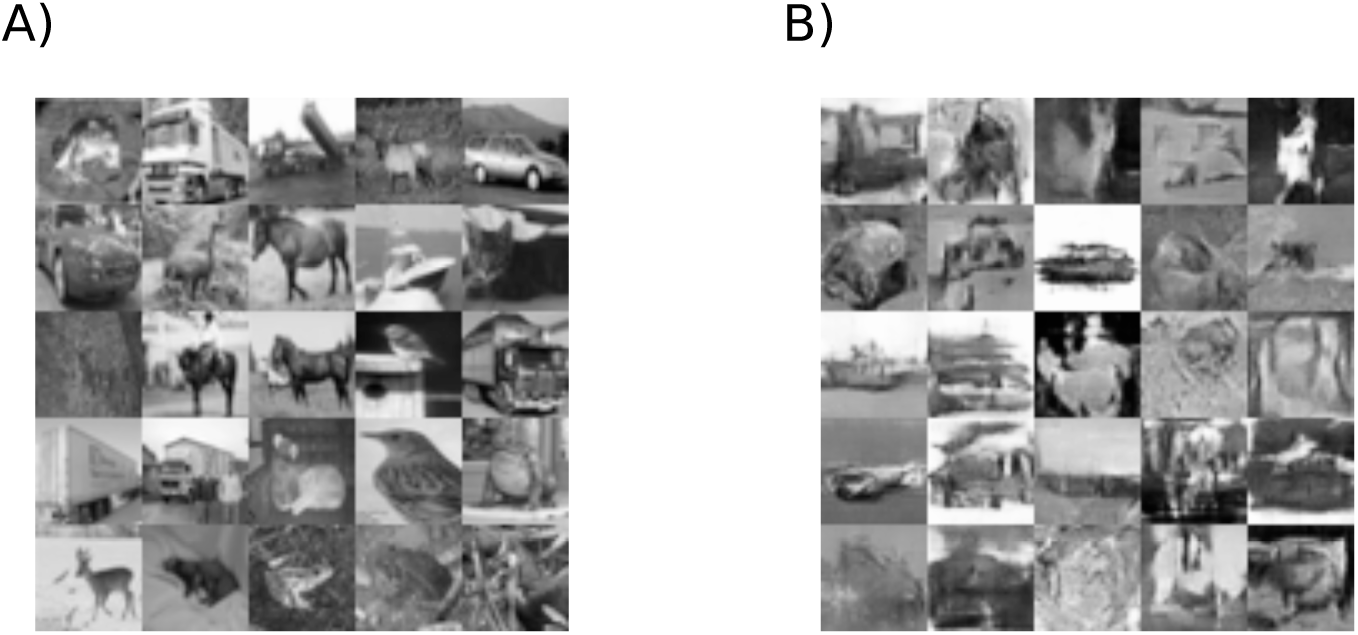
Example training images and samples. **A**. Training images from the CIFAR10 database used to train the GAN model. **B**. Samples generated from the trained GAN. Similar image samples were used in the experiment.

**Figure 2:**
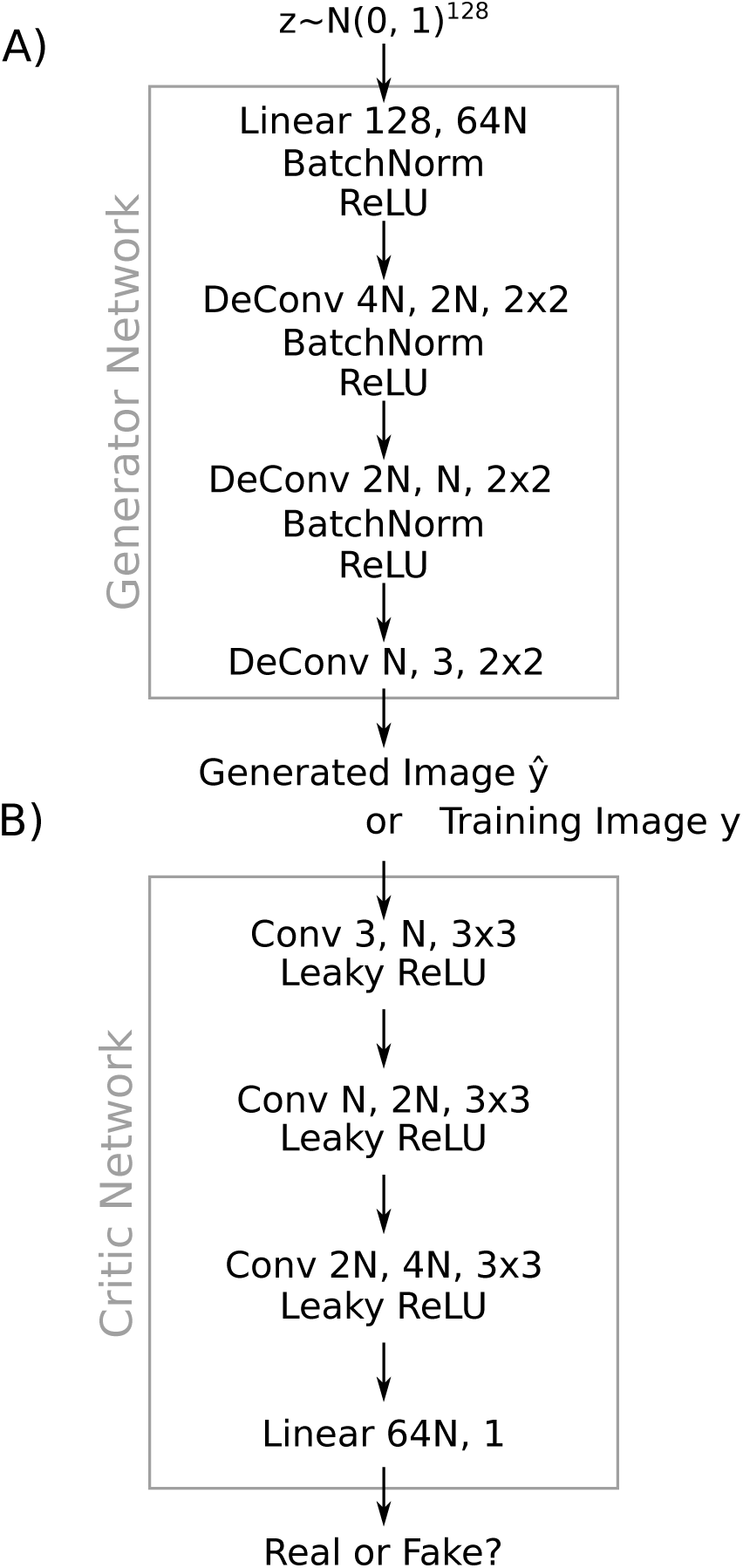
Architecture of the generative adversarial network. **A**. Architecture of the generator network. Information flow is from top to bottom, network layers are “Linear *k*, *m*”: Affine transformation from *k* features to *m* features, “Conv *k, m, n* × *n*”: Convolutional layer from *k* channels to m channels using a kernel size of *n* × *n*, “DeConv *k*, *m*, *n* × *n*”: like convolution but up-upsampling before the convolution to increase spatial resolution and image size, “BatchNorm”: Batch normalization (Ioffe & Szegedy, 2015), “ReLU”: rectified linear unit ReLU(*x*) = max(0, *x*) (Glorot et al., 2011). The generator network maps a sample *z* from an isotropic 128 dimensional Gaussian to a 32 × 32 pixel colour image. **B**. Architecture of the critic network. Architecture components not used in A. are “Leaky ReLU” (He et al., 2015). The critic network receives as input either an image 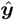 generated by the generator network or a real training image ***y***, and it decides if the input image is real or not.

Networks with different numbers of hidden states (parameter *N* in Figure 2) were trained for 200 000 epochs using an ADAM optimizer (Kingma & Ba, 2015) with learning rate 10^−4^ and *β*_0_ = 0, *β*_1_, = 0.9. Specifically, we trained networks with *N* = 40, 50, 60, 64, 70, 80, 90,128 (see Figure 2). Wasserstein-2 error (Arjovsky et al., 2017) on a validation set (the CIFAR10 test dataset) was lowest with *N* = 90 in agreement with visual inspection of sample quality, so we chose a network with *N* = 90 for all remaining analyses. Example images generated from this final network are shown in Figure 1B.

#### Observers

Seven observers participated in the experiment. Two of them were authors, the remaining five were students from various labs at the Centre for Vision Research at York University, Toronto, Ontario. One additional observer participated in the first session, but withdrew from the experiment afterwards and their data was excluded from the analysis. All observers reported normal or corrected-to-normal vision. Prior to participation, all observers provided written informed consent to participate and all procedures were approved by the York University Ethics Board.

#### Procedure

Each individual observer judged image samples from the GAN in a spatial two-alternative forced choice match-to-sample task (see Figure 3). On every trial, the observer saw three images; a target image at the center, one comparison stimulus on the left and another on the right. One of the comparison stimuli was a perturbed version of the target image generated from the GAN, while the other was an equally perturbed version of a seperate image generated from the GAN. The observer was required to indicate which of the two comparison images matched the central target image by pressing a corresponding button on a computer keyboard (left arrow key if the left comparison stimulus matched, right arrow key if the right comparison stimulus matched). Stimuli were presented for up to 6s or until the observer made a response, resulting in practically unlimited viewing time. Before each trial, a fixation point appeared on the screen for 500ms. Each observer performed 5 sessions and each session consisted of 80 trials for each noise level and type, resulting in a total of 1300 trials per observer (except O2, who performed 1437 trials).

**Figure 3:**
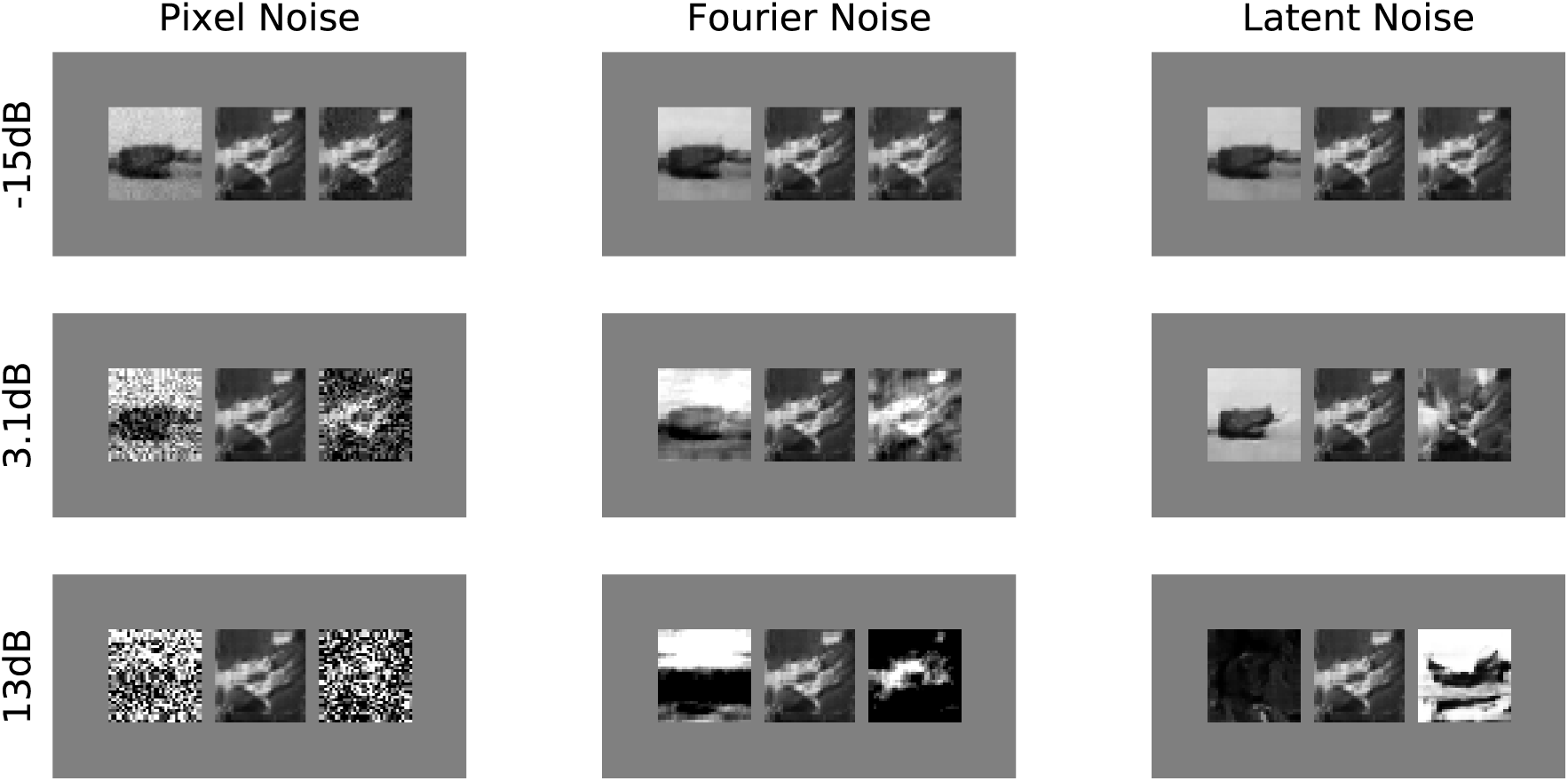
Design of the image matching experiment. On every trial, the observer saw three images; a target image at the center, one comparison stimulus on the left and another on the right. Comparison stimuli were sampled randomly from the GAN and perturbed by different types of noise. Pixel noise (left column) was added as independent Gaussian noise to every pixel, Fourier noise (middle column) with the same power spectrum as the original image, but with random phases, was added to the original image, Latent noise (right column) was independent Gaussian noise applied in the latent space of the GAN that was used to generate the images. The top row shows example experimental displays with low noise, the central row shows example experimental displays close to the critical noise level for latent stimuli, the bottom row shows example experimental displays with high noise (amounts given on left). For illustration, the left stimulus is a random perturbed stimulus and the right stimulus is the perturbed target in every example display. During the experiment, the identities and locations of the comparison stimuli were randomized.

#### Stimuli

All stimuli were samples from a GAN, converted to gray scale by averaging the red, green and blue channels of the sample image. The target stimulus was always noise-free, while the two comparison stimuli were perturbed by one of three noise types (see Figure 3). Pixel noise was constructed by adding independent Gaussian noise to each pixel of the respective image. Fourier noise was constructed in the Fourier domain by replacing the image’s phase component by random numbers. This resulted in an image with the same power spectrum as the source image but with completely random phases. A Fourier-perturbed image was constructed by adding a multiple of this power-spectrum-matched noise to the source image. Perturbations of the latent vectors of a GAN are closely correlated to pixel space perturbations (*r* = 0.82, *p* < 10^−60^), but they are not exactly the same. Therefore, we constructed latent noise by manipulating the latent vector ***z*** from which an image was generated. To generate perturbed images with a predefined difference in pixel space, we started by adding independent Gaussian noise *ζ* to ***z*** and determining the corresponding image *G*(***z*** + *ζ*). We then used gradient descent on

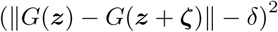
to adjust *ζ* such that the final difference between the target and the perturbed target had a predefined pixel space difference of *δ*.

Stimuli were presented on a medium gray background (54.1 cd/m^2^) on a Sony Triniton Multiscan G520 CRT monitor in a dimly illuminated room. The monitor was carefully linearized using a Minolta LS-100 photometer. Maximum stimulus luminance was 106.9 cd/m^2^, minimum stimulus luminance was 1.39 cd/m^2^. If the nominal stimulus luminance exceeded that range, it was clipped (for subsequent analyses, we also used the clipped stimuli). On every frame, the stimuli were re-rendered using random dithering to generate a quasi-continuous luminance resolution (Allard & Faubert, 2008). At a viewing distance of approximately 87cm, each stimulus image subtended approximately 0.65 degrees of visual angle and were separated by approximately 0.13 degrees of visual angle. One pixel subtended approximately 0.02 degrees of visual angle.

#### Data analysis

For every observer, we estimated a psychometric function parameterised as

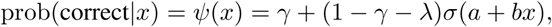
where *γ* = 0.5 is the probability to guess the stimulus correctly by chance, *λ* is the lapse probability, *σ* is the logistic function and *a* and *b* govern the offset and the slope of the psychometric function. Here, *x* is the root-mean-square level of noise applied to perturb the respective images in dB relative to the screen’s background luminance. However, we note that different ways of scaling the noise (other than dB) did not impact our main results. We adopted a Bayesian perspective on estimation of the psychometric function (Fründ et al., 2011) and used weak priors *λ* ~ Beta(1.5,20), *a* ~ *N*(0,100), *b* ~ *N*(0,100), where *a* and *b* are expressed on the dB scale of the noise. Mean *a posteriori* estimates of the critical noise level *x_c_* at which *ψ*(*x*) = 0.75 and the slope of the psychometric function at *x_c_* were obtained using numerical integration of the posterior (Schütt et al., 2016).

To understand how the structure of the GAN’s latent space determined image matching performance, we re-analysed data from the latent condition. For this re-analysis, we assumed that observers would pick the perturbed stimulus that is closer to the target with respect to some distance measure. More specifically, let ***t*** denote the noise-free target stimulus and 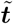 and 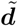 denote the perturbed target and distractor stimuli respectively. If an observer is picking the stimulus that is closer to the target, then 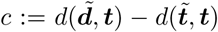 should show a positive correlation with the observer’s trial by trial response accuracy. Here, *d* is a suitably defined distance measure. We used either the Euclidean distance 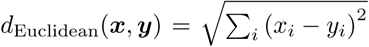, the radial distance 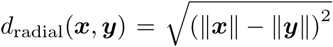, or the cosine distance 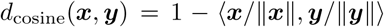, where ||*x*|| denotes the Euclidean norm of a vector ***x*** and 〈***x***, ***y***〉 denotes the scalar product of vectors ***x*** and ***y***. These distances were applied in either the GAN’s latent space or directly in pixel space, after concatenating the respective stimulus’ pixel intensities into one long vector. We then determined receiver operating curves (ROC) for predicting correct vs incorrect responses based on *c*. The area under the ROC is a measure for how well the respective distance measure predicts the observer’s trial by trial responses (Green & Swets, 1966). To test if the area under the curve (AUC) was significantly different from chance, we performed a permutation test randomly re-shuffling the correct/incorrect labels 1000 times and taking the 95-th percentile of the resulting distribution as the critical value. We also used permutation tests to determine if the AUC for two different distance measures was significantly different. For the pairwise comparisons, there are 128 possible re-assignments of AUC values to the two conditions, and we computed all of them. The *p*-values for these post-hoc comparisons were corrected for multiple comparisons to control for inflation of the false discovery rate (Benjamini & Hochberg, 1995).

To gain insight into the image features that determined the observers’ responses, we applied the ROC analysis to a number of image features as well. Firstly, we calculated the luminance histogram (50 bins) for each image and calculated the distance difference *c* between luminance histograms of the respective images. Distance measures are computed over vectors of length 50, and each entry denotes the bin count. Secondly, to determine local dominant orientation at each pixel we first filtered the image with horizontal and vertical Scharr filters (Scharr, 2000) as implemented in scikit-image (van der Walt et al., 2014) giving local horizontal structure *h* and vertical structure *v*. The local orientation *ϕ* was extracted from these two responses such that *h* = *r* cos(*ϕ*) and *v* = *r* sin(*ϕ*), where 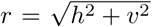. We then determined the histogram (3 bins) of the local orientations across the image and calculated *c* as the distance difference between these orientation histograms. As a third feature, we calculated the edge densities of the two images, using the canny edge detector from scikit-image with a standard deviation of 2 pixels and calculating the fraction of pixels labeled as edges by this algorithm. As a fourth feature we determined the slope of the power spectrum in double logarithmic coordinates.

Finally, we used a standard method for image segmentation (Felsenzwalb & Huttenlocher, 2004) to calculate segmentations of the images ***t***, 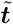 and 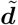. We used the method implemented in scikit-image (van der Walt et al., 2014). Briefly, this algorithm iteratively merges neighbouring pixels or pixel groups if the differences across their borders are small compared to the differences within them. Each segmentation consists of a number of discrete labels assigned to the pixels of the original image. If two pixels belong to the same segmented region, the two labels associated with them should be the same. However, different segmentations may assign different labels to the same region. Thus, two segmentations ***s*** and 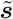 would be similar, if for many pairs (*i, j*) of pixels ***s****_i_* = ***s****_j_* implies 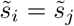. When calculating the distance between two segmentations ***s*** and 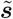, we therefore count the number of pixel pairs for which ***s****_i_* = ***s****_j_* and 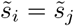 and we normalize by the number of pixel pairs that are assigned to the same region by at least one of the two segmentations. This is captured by the distance measure

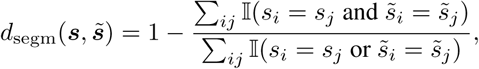
where 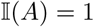 if the expression *A* is true and 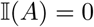 otherwise and the sums go over all pairs of pixels *i*, *j*. If the two segmentations define exactly the same regions (but possibly with different labels), *d*_segm_ will be 0, if the two segmentations are completely different, in the sense that one has only one region (the entire image) and the other assigns each pixel to its own region, then *d*_segm_ will be 1.

### Results

#### Lower tolerance for noise applied within an approximation of the natural image manifold

In the first experiment, seven observers were required to judge which one of two noisy comparison images matched a centrally presented target stimulus (see Figure 3). Figure 4A shows psychometric functions for one example observer as a function of noise level. In general, higher noise levels were associated with less correct responses. The observer’s performance, indicated by level of response correctness, was least affected by noise that was applied independently to each pixel (critical noise level with 75% correct performance at 14.8 ± 1.43dB, posterior mean and standard deviation, blue dots and line in Figure 4A). Observer performance was more affected by noise that was applied in the pixel domain but matched the power spectrum of the original image (critical noise level at 7.31 ± 0.54dB, green dots and line in Figure 4A). Finally, the level of response correctness was most affected by noise that was applied in the latent space of the GAN and thus stayed within an approximation of the manifold of natural images (critical noise level at 1.60 ± 0.60 dB, red dots and line in Figure 4A). Thus, this observer’s level of response correctness was most affected by noise applied within a model of the manifold of natural images. Furthermore, the psychometric function for this observer was considerably steeper in the latent noise condition (slope at critical noise level was −0.11 ± 0.09/dB) than in the other two conditions (slopes at critical noise level were −0.015 ± 0.006/dB for pixel noise and −0.013 ± 0.0015/dB for Fourier noise), suggesting that the observer was relatively insensitive to low amplitude latent noise and then abruptly became unable to do the task, consistent with the idea that for latent noise above a certain level a categorical change happens, while noise in the pixel domain results in a more gradual decrease in image quality.

On average across all 7 observers, the critical noise level was highest for independent pixel noise (8.91±1.22dB, mean±s.e.m.) and it was comparable for Fourier noise (6.77±0.83dB, paired *t*-test pixel vs. Fourier noise: *t*(6) = −1.59, n.s.) and decreased significantly for latent noise (3.47±0.36dB, paired *t*-test pixel vs. latent noise: *t*(6) = 3.59, *p* = 0.011, Fourier vs. latent noise: *t*(6) = 3.89, *p* = 0.0080) respectively (see Figure 4B). Thus overall, observers were most affected by noise that was approximately applied within the manifold of natural images by perturbing the GAN’s latent representation of the stimulus. We verified that this result also held for every individual observer. We further found that psychometric functions tended to fall off more steeply when noise was applied in the GAN’s latent space (average slope at critical noise level for latent noise was −0.066±0.0084/dB, see Figure 4C) than when noise was applied in pixel space (average slope at critical noise level for pixel noise was −0.024±0.0041/dB, for Fourier noise −0.017±0.0027/dB), replicating the observations from Figure 4A.

**Figure 4:**
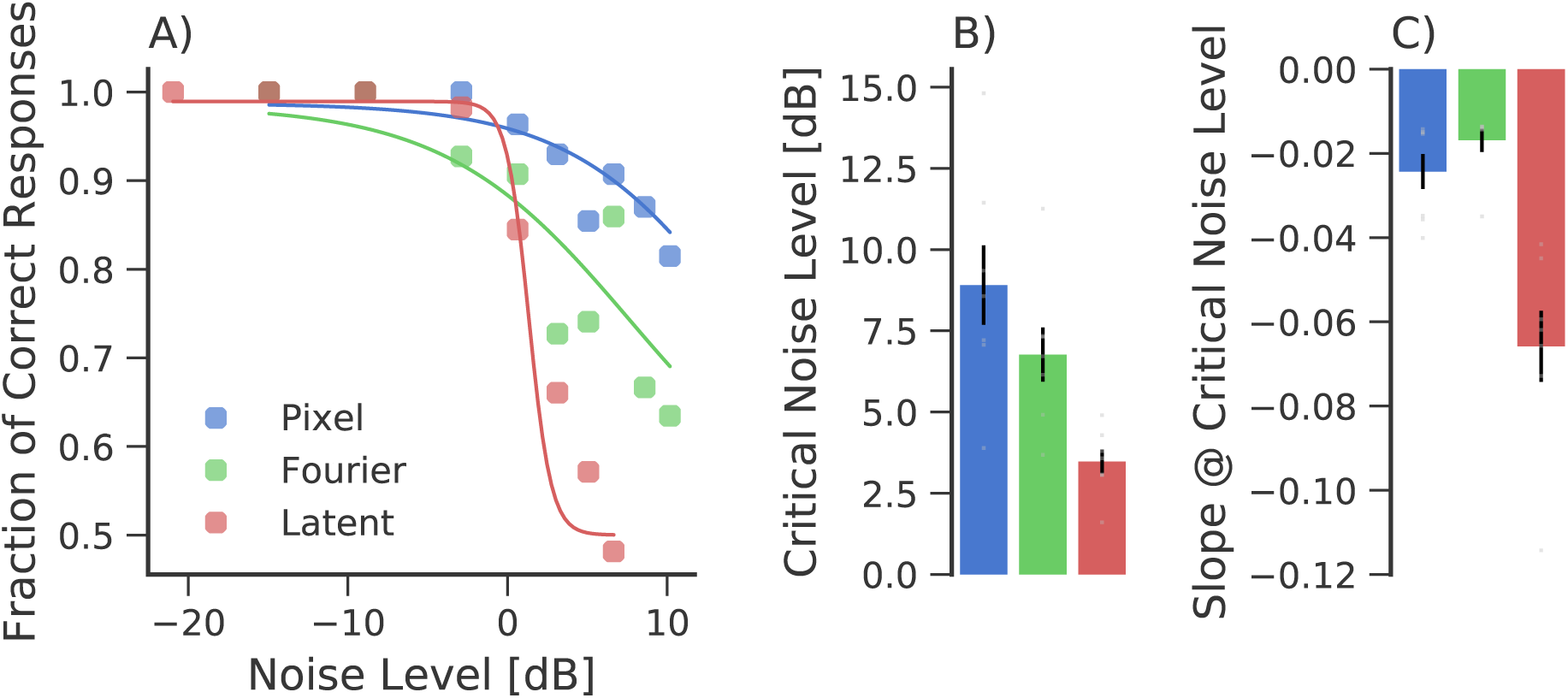
Higher sensitivity to noise perturbations applied within the manifold recovered by a GAN. **A**. Psychometric function for one example observer. Each dot represents between 50 and 60 trials, solid lines are mean a-posteriori estimates of the psychometric function (see Methods). All noise levels were quantified as root mean square difference to the target in pixel space and were normalized to the background luminance. **B**. Average critical noise levels corresponding to 75% correct performance. Height of the bars denotes the mean across 7 observers, error bars indicate s.e.m. across observers. Light gray dots indicate results for individual observers. Colors are the same as in part A. **C**. Average slope of the psychometric function at the critical noise level. Height of the bars denotes the mean across 7 observers, error bars indicate s.e.m. across observers. Light gray dots indicate results for individual observers. Colors are the same as in part A.

To summarize, we found that, in general, noise that was approximately applied within the manifold of natural images was more effective at interrupting observers’ performance than noise applied outside of the manifold of natural images in pixel space. We performed two additional sets of analysis to determine (i) if latent space or pixel space image differences were more predictive of observer’s trial by trial behaviour and (ii) which image features were responsible for the decline in performance in the latent noise condition.

#### Distance in latent space correlates with image matching performance

We modeled observer behaviour in the image matching task, by assuming that on every trial, the observer picks the comparison stimulus that appears closer to the target with respect to some appropriate distance measure. We looked at three different candidate distance measures and applied them both in the GAN’s latent space and directly in pixel space. To evaluate the relevance of each distance measure, we non-parametrically calculated area under the receiver operating curve (AUC) values for discrimination between correct and incorrect trials.

Figure 5A shows average AUC-values for different hypothetical distance measures applied either in latent space or in pixel space. The simplest distance measure is the Euclidean distance, marked by the blue bars in Figure 5A (darker blue for Euclidean distance in latent space, lighter blue for Euclidean distance in pixel space). Note that Euclidean distance in pixel space is equivalent to the standard deviation of the difference image between stimuli that was used as a unified measure of perturbation strength in Figure 4 (without the transformation to dB). Euclidean distance in latent space was at least as predictive as Euclidean distance in pixel space (latent space: 0.83±0.0092, average AUC ± s.e.m., pixel space: 0.82±0.014, permutation test *p* = 0.17).

**Figure 5:**
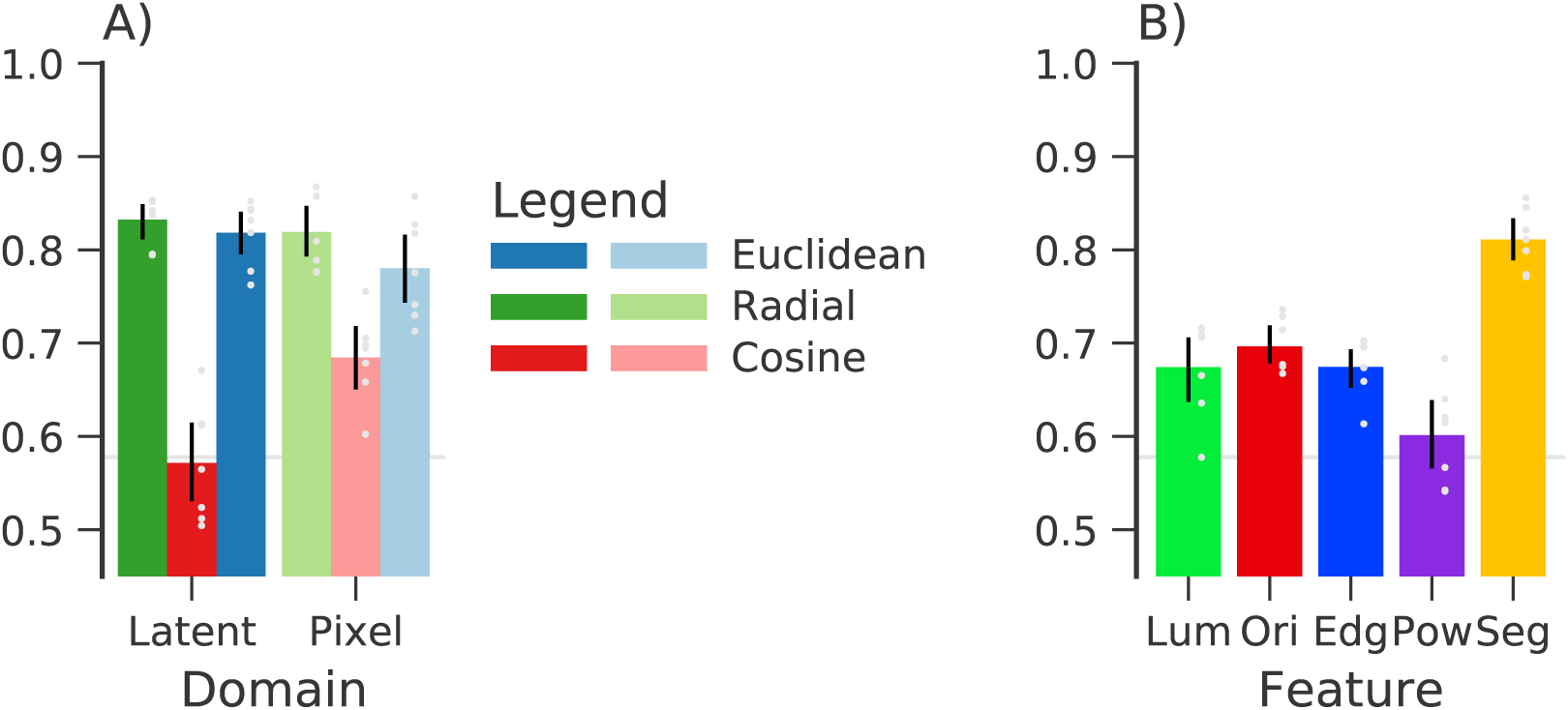
Explaining trial by trial performance. **A**. Area under the ROC curve for different hypothetical distances applied in latent space or pixel space. Distance measures applied in the GAN’s latent space, corresponded to distances within a model of the manifold of natural images (darker colors). Color hue codes for the type of distance measure used. Error bars indicate 95% bootstrap confidence intervals. The light gray line marks the average critical value for chance performance. **B**. Same as A. but for image features. “Lum” corresponds to comparisons of luminance histograms, “Ori” corresponds to comparisons of orientation histograms, “Edg” corresponds to edge density, “Pow” corresponds to the slope of the image’s power spectrum, and “Seg” corresponds to comparisons of image segmentations (see Methods for detail).

We performed the same analysis using the difference between the norms of the either the latent vector or the pixel vector (green bars in Figure 5A). In pixel space, radial distance is equivalent to an observer who simply compares the RMS contrast of the images. In contrast to the Euclidean distance, this observer would *first* take the standard deviation of the target and flanker images and then compare differences in standard deviations. In latent space, the norm of the latent vector seems to be related to contrast as well, but the relationship is more complex. Radial distance receives considerably lower AUC than Euclidean distance in both latent and pixel space and was a much less reliable predictor of trial by trial performance. Radial distance in latent space was significantly less predictive than radial distance in pixel space (latent space: 0.57±0.022, pixel space: 0.68±0.016, permutation test *p* < 0.05 corrected) and for four out of 7 observers, radial distance in latent space did not predict trial by trial choices significantly better than chance.

Finally, we analyzed how well cosine distance explained observers’ responses (red bars in Figure 5A). Cosine distance is interesting for two reasons. Firstly, cosine distance is equivalent to Euclidean distance except for the influence of the radial component.

Secondly, cosine distance is closely related to correlation and an observer who relies on cosine distance essentially uses the target stimulus as a template and computes the correlation with each of the comparison stimuli to pick the stimulus that correlates best with that template^1^. Cosine distance applied in latent space was a better predictor than if it was applied in pixel space (latent space: 0.82±0.012, pixel space: 0.78±0.019, permutation test *p* = 0.05 corrected).

In the appendix, we replicate these results using an analysis based on logistic regression.

Taken together, the results of image features and hypothetical distance measures suggest that the latent space of GANs captures processing beyond simple contrast differences. Specifically, trial-by-trial responses were most accurately explained by differences in image segmentations rather than low-level features such as luminance, orientation, or edge density.

#### Distortions of mid-level features explain trial by trial performance

In order to determine which image features were responsible for the decline in performance with perturbations in the latent space of GANs, we applied the same analysis to different image features (see Figure 5B).

We found that differences in the luminance distribution of the images are clearly predictive of trial by trial behaviour (average AUC: 0.67±0.018, AUC was larger than 95-th percentile of the null distribution in all seven observers). Yet, other features, such as the difference in local orientation (average AUC: 0.70±0.010) or differences in the images edge density (average AUC: 0.67±0.011) were equally good predictors of the observers’ trial by trial behaviour (permutation test not significant after correction for multiple comparisons).

One of the quantities that might have been relevant for our observers is the slope of the power spectrum (see for example Alam et al., 2014). We evaluated to what extent this feature contributed to our observers’ decisions and found that it is largely irrelevant at explaining the observers’ trial by trial behaviour: In three out of seven observers the AUC of this feature was not significantly different from chance performance and the average AUC for the slope of the images’ power spectrum was significantly less than that for any other feature we studied.

In order to quantify how well differences in the mid-level structure of the perturbed images could explain trial by trial responses, we calculated segmentations of all images using a standard segmentation algorithm (Felsenzwalb & Huttenlocher, 2004). As this feature is less common than the other image features, we show the distribution of this image feature in the appendix. Although we do not believe that humans necessarily segment images using graph based optimization as the algorithm does, we believe that this approach provides at least a coarse approximation to the mid-level structure of the images. Differences in segmentation were considerably more predictive than differences in any other feature distribution (mean AUC for segmentation 0.82±0.011, permutation test *p* < 0.05 corrected). In fact, differences in segmentation were about as predictive of trial by trial behaviour as cosine distance in latent space (permutation test *p* = 0.60), suggesting that indeed distortions of the images’ mid-level structure might be responsible for the decline in image matching performance when noise was constrained to stay within the recovered manifold of natural images by applying it in the GAN’s latent space.

## Experiment 2: Sensitivity to directions in the recovered natural image manifold

In Experiment 1, we found that human observers are particularly sensitive to image perturbations that stay within an approximation of the manifold of natural images. This was achieved by perturbing stimuli along a parameterization of the manifold of natural images as recovered by a GAN. We wondered if observers were also sensitive to other aspects of this parameterization, such as for example direction. We therefore asked observers to discriminate between videos that were created by walking along either straight paths in latent space (i.e. which contained no change in direction) or paths that had a sudden turn (i.e. which contained a change in direction).

### Method

#### Observers

Five observers participated in the second experiment and the control condition for the second experiment. All five of them had participated in the first experiment as well. The procedures were approved by the Ethics Board of York University, ON, Canada and observers provided informed consent before participating. One observer (O3) accidentally did one session with incorrect angle logging. As the correct angles could not be recovered, we decided to exclude the corresponding trials from the analysis.

#### Procedure

The experiment was a single interval design. On every trial, the observers either saw a video without a turn in latent space direction or a video that contained a turn in latent space direction and they had to decide if the presented video contained a turn or not. The probabilities for turn and no-turn videos were each 0.5. In order to avoid bias about the image features that would indicate a turn in latent space, observers were instructed that there would be two classes of videos and that we were not able to describe the difference unambiguously in words. Instead, each observer first saw 100 trials with path angles of 90° (vs straight) and received trial by trial feedback about their performance. After that, each observer performed two blocks of 100 trials for each path angle for the main experiment. For each block, the size of the possible path angle was kept constant. To allow the observers to calibrate their decision criterion to the size of turns presented in the respective block, we provided trial by trial feedback during the first 20 trials of each block and only analyzed the remaining 80 trials. Thus in total, we analyzed 160 trials per observer per path angle.

#### Stimuli

Each video consisted of 60 frames. The first frame corresponded to a random point on the sphere with radius 10. Successive frames were then constructed by taking steps of norm 0.5 in a random direction tangential to the sphere with radius 10 (see Figure 6A). Straight paths were constructed by simply taking 60 successive steps in the same direction. Paths with a turn were constructed by changing the direction of steps by an angle *α* of 15, 30, 60, or 90 degrees after the first 30 frames. We will refer to this angle as the “path angle” in the following. At a frame rate of 60Hz, each video had a duration of 1s and if the video contained a turn in latent space, that turn happened after 500ms. Otherwise, the setup for experiment 2 was the same as in experiment 1.

**Figure 6:**
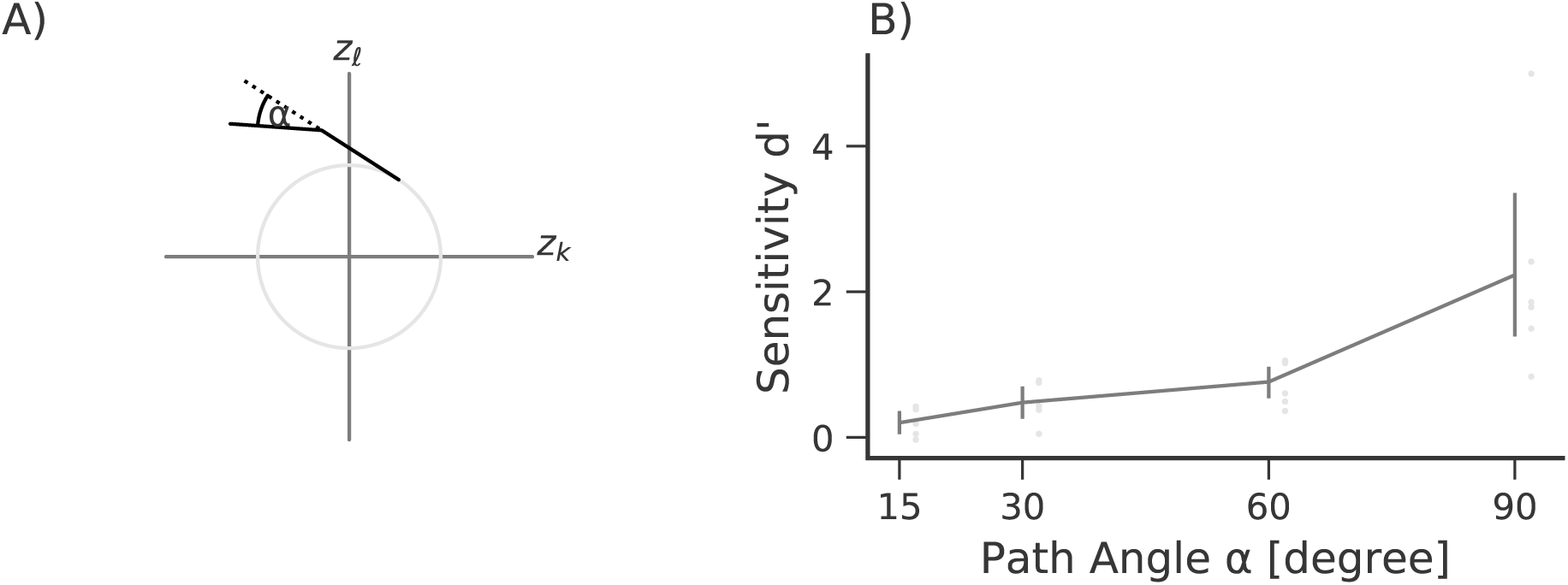
Observers can detect turns in paths along the manifold of natural images as recovered by a GAN. **A**. Illustration of stimulus construction. Two latent axes ***z****_ℓ_* and ***z****_k_* are shown for illustration, the actual latent space had a dimension of 128. The light gray circle marks the sphere of radius 10 in latent space. Shown are a straight path (dotted line, standard stimulus in the experiment) and a path that turns half way at an angle *α* (solid line, target stimulus in the experiment). **B**. Average sensitivity for detecting a turn in an image sequence along a path along the modeled manifold. Error bars mark 95% confidence intervals. Light gray dots mark individual observer results.

For the control experiment, additional videos were constructed for paths through pixel space, through Fourier space, and through latent space. Videos for paths through pixel space were constructed in the same way as described above with the only difference that the respective vectors were sampled from an isotropic Gaussian. Videos for paths through Fourier space were similar to videos constructed in pixel space, but were additionally filtered to an approximate 1/*f* power spectrum by multiplying their Fourier transform with 1/(0.1+*f*). The exact size of the rms difference between successive video frames varied somewhat from trial to trial, but there were no statistically significant differences between frame by frame differences in pixel space, in Fourier space, or in latent space (*p* > 0.1).

#### Data analysis

To evaluate observers’ sensitivity to path angles, we calculated *d*′ at each path angle. For single observers, confidence intervals for *d*′ were determined by bootstrap with 1000 samples.

### Results

Figure 6B shows average discrimination performance as a function of path angle. Not surprisingly, observers were best at detecting turns of 90 degrees in latent space (average *d*′ = 2.23 ± 0.59). However, even for path angles of 15 degrees, the smallest path angles tested, three out of five observers performed above chance (average *d*′ = 0.20 ± 0.090, one-sided *t*-test against zero: *t*(4) = 2.25, *p* = 0.0436). This indicates that even small changes in latent space direction were detected by the observers.

Overall, observers were remarkably good at discriminating between different directions taken by movies in latent space. In fact half of the observers responded correctly in more than 80% of the trials (average 78.1±2.56%, mean ± s.e.m.) at the largest turn.

Is sensitivity to these videos really specific for the latent space of a generative adversarial network? To control for the possibility that observers could simply detect any abrupt change in continuously morphed images, we performed a control experiment in which we used only 90-degree turns but varied the space in which the videos corresponded to straight paths. We found that observers were insensitive to turns in pixel space paths (average *d*′ = −0.16 ± 0.29 mean ± s.e.m) and were moderately sensitive to turns in Fourier space paths (average *d*′ = 1.09 ± 0.24). However, sensitivity to turns in paths through the GAN’s latent space was highest (average *d*′ = 1.89 ± 0.20). The pairwise comparisons between performance for pixel space vs Fourier space paths (*p* < 0.05, permutation test) and performance for Fourier space paths vs latent space paths (*p* < 0.05, permutation test) were both statistically significant with the observed configuration having the largest difference of all permutations. This confirms that sensitivity to directions in latent space is a specific property of the representation recovered by generative adversarial networks.

## General Discussion

We found that observers are sensitive to image manipulations that are restricted to remain within an approximation of the manifold of natural images. Errors made within the manifold of natural images recovered by a GAN seem to be related to changes in the configural structure of the images more than to changes in local image features. We further find that observers are sensitive not only to noise within the model of the manifold, but also to more subtle aspects of paths along the model of the manifold (see Experiment 2).

### Global vs local image models

Our results seem to contradict reports which find that observers are very sensitive to deviations from naturalness (Gerhard et al., 2013): How would observers be equally sensitive to changes within the manifold of natural images and deviations away from that manifold? We find that sensitivity to perturbations within an approximation of the manifold of natural images can be well predicted by the part and object structure of the image that is captured by standard segmentation algorithms (see Figure 5), while deviations from naturalness might be more related to the fine structure of local statistics. This is also visible in Figure 1: The two image classes in parts A. and B. appear very similar at first glance and only upon closer inspection does it become clear that the examples in part B. are mostly meaningless. We believe that this point could be taken both as an advantage and a disadvantage. Clearly, the samples in Figure 1B. do not match every aspect of the natural images in Figure 1A. (although better matches can be achieved if the GAN is restricted to more narrow classes of objects, see for example Radford et al., 2016; Gulrajani et al., 2017). However, the images capture a lot of the global and highly non-stationary properties of natural images, that texture models based on stationarity do not capture (Portilla & Simoncelli, 2000; Gatys et al., 2015). We therefore believe that our approach is complementary to the local approach taken by studies that investigated texture processing (e.g. Freeman & Simoncelli, 2011; Gerhard et al., 2013; Wallis et al., 2016, 2017)

**Figure 7:**
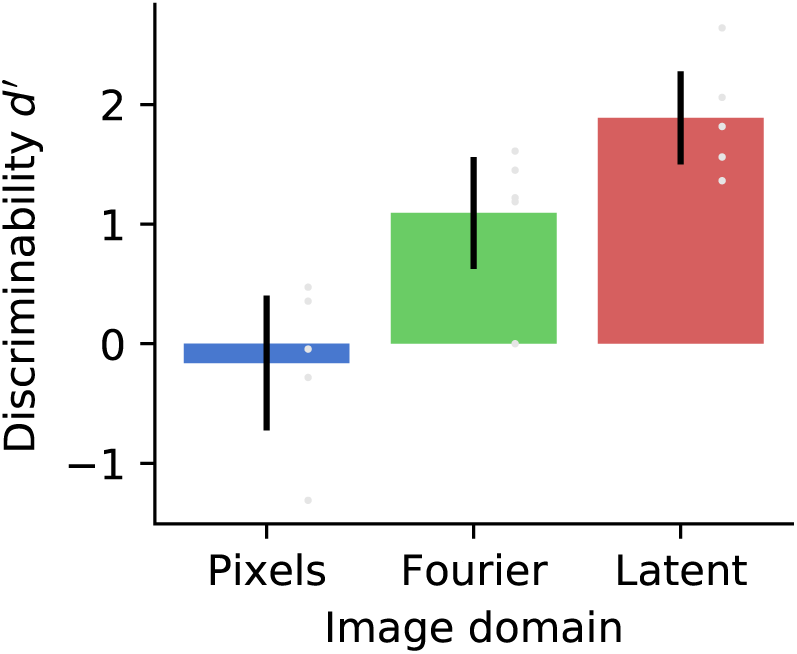
Turns in other image spaces are harder to detect. Average sensitivity for detecting a 90-degree turn in an image sequence along a path through pixel space, through Fourier space, and through latent space. Error bars mark 95% confidence intervals and gray dots mark performance of individual observers.

### Small images

The images employed here are relatively small. Our images were only 32×32 pixels in size. In contrast, Wallis et al. (2017) used image patches that were 128 × 128 pixels to compare between texture images created from a deep neural network model and real photographs of textures. Other studies have used a range of images sizes (Alam et al., 2014; Sebastian et al., 2017; Bex, 2010, in increasing order of image size), but our images are closer to the range of image sizes used as patches of images (Gerhard et al., 2013) rather than entire images. However, training generative adversarial networks on larger images with similar image variability as the CIFAR10 network currently typically requires training class conditional networks as for example done by Miyato et al. (2018) when training on the entire ImageNet dataset (Russakovsky et al., 2015). Although this would have in principle been possible in this study as well, it would have implied that separate manifolds would be used for different classes and it would have made interpretation of our results considerably more complex. We therefore decided to restrict ourselves to smaller images.

### Dataset bias

Many publicly available data bases appear to be systematically biased (Wichmann et al., 2010): The images in these databases are pre-segmented in the sense that a photographer selected a viewpoint that they considered particularly “interesting” or in that they selected which objects to put in focus. Although pictures from these databases may appear natural, conclusions drawn from these data bases may be misleading (Wichmann et al., 2010). The example pictures in Figure 1A. clearly show such photographer bias. In every one of these examples, the perspective is clearly focused on one specific object, while typical natural scenes often contain multiple objects and a lot of not explicitly defined random texture (i.e. background). In general, the extent to which such bias would influence our results depends on the extent to which the biased pixel-by-pixel statistics in these images determine responses in our task. In the section “Distortions of mid-level features explain trial by trial performance” we show that the main determinant of responses in our experiments was the segmentation structure of our images. Although these might also be influenced by photographer bias, we assume that the impact is more subtle. Furthermore, eye movements tend to center objects on or close to the fovea (Kayser et al., 2006; ’t Hart et al., 2013), which might lead to similar global effects as a photographer focussing the image on selected objects. However, the extent to which photographer bias impacts the image representation recovered by GANs is still an open question.

### Non-object images

Upon closer inspection, the examples in Figure 1B. do not look exactly like real objects. Although each one of the examples can clearly be segmented into foreground and background, it is not always possible to actually name the foreground objects in Figure 1B., while this seems to be easier for the training examples in Figure 1A., despite the relatively low resolution of the images. Thus, the samples from the GAN used in the present study would probably be easy to discriminate from real images if they were directly compared to real images (Gerhard et al., 2013; Wallis et al., 2017). It should thus be noted that the image representation learned by the GAN used in this study is only an approximation to the manifold of natural images (note however, that other studies training on the CIFAR10 dataset show samples of similar quality, for example Gulrajani et al., 2017; Roth et al., 2017). Although image manipulations in this approximation appear to be convincing if the GAN has been trained on more restricted sets of training images (see for example Zhu et al., 2016), this can not be guaranteed for all the stimuli used in this experiment. To this date, it is unclear how exactly the GAN samples used in this study match the perceived properties of the training data (even though the training data themselves might be biased, see Section “Dataset bias”).

We believe however, that natural-appearing non-objects are still an interesting class of stimuli. For example, Huth et al. (2012) report that semantic content is important for shaping the response properties of large parts of anterior visual cortex, suggesting that many areas that are traditionally thought of as visual, are also semantic areas. Attempts to further test this claim have been restricted to correlational approaches (Khaligh-Razavi & Kriegeskorte, 2014), partly because it is difficult to generate “non-object” stimuli that otherwise fully match the properties of object stimuli (see also Fründ et al., 2008). Comparing responses to training images with objects (Figure 1A) to images generated from a GAN but without the full semantic information (Figure 1B) could help resolve this point.

### Conclusion

In this study, we explored the potential of studying vision within an approximate manifold of natural images. To do so, we employed a generative adversarial network to constrain perturbations to remain within the manifold of natural images and we find that observers are remarkably sensitive to image manipulations that are constrained in this way. We observe that perturbations within a model of the manifold of natural images tend to disrupt more global image structures such as figure-ground segmentation structure. This might prove useful in future studies that investigate such processes under more naturalistic conditions. The fact that GANs provide an approximate parametrization to the manifold of natural images encourages further use of these powerful image models to study vision under complex naturalistic stimulus conditions.

All code and data for this manuscript is available online at http://doi.org/10.5281/zenodo.1308853.

## Appendix

## Distribution of segmentation distances

Although the distributions for most image features analyzed in Experiment 1 is relatively intuitive, this might not be the case for the segmentation distances. Figure 8 shows the distribution of segmentation distances *d*_segm_ across all trials.

**Figure 8:**
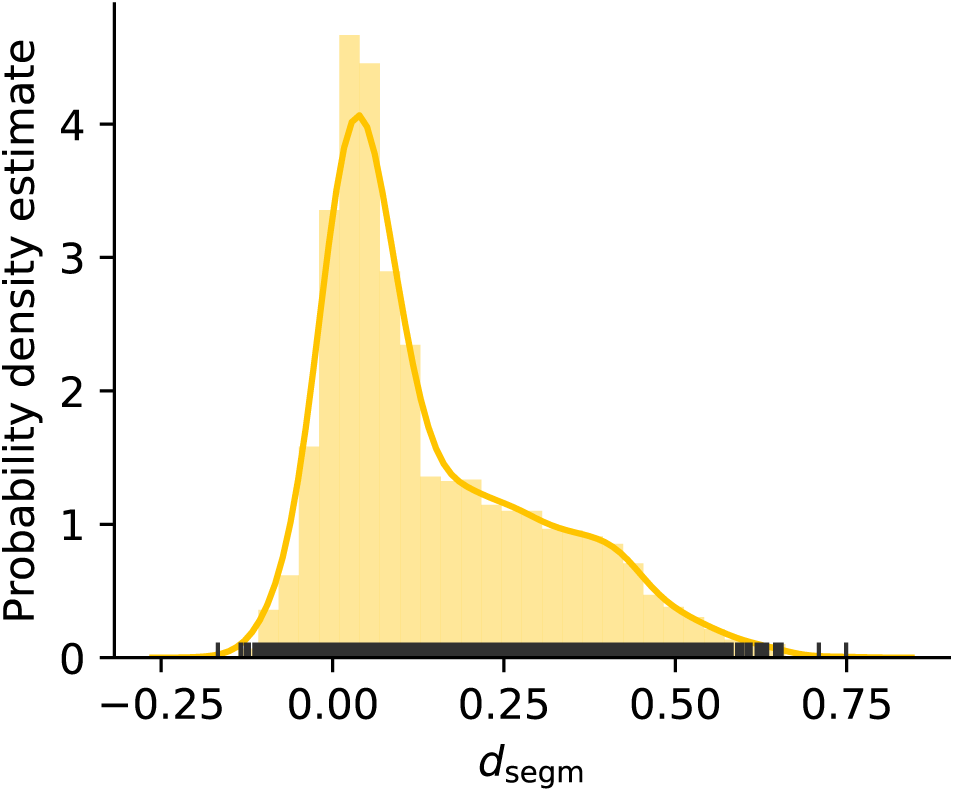
Distribution of segmentation distances across all trials. The yellow line is a kernel density estimate of the distribution, the bars show a histogram of the distribution. Kernel band width has been determined by Scott’s rule (Scott, 1992), bin size has been determined by Freedman-Diaconis (Freedman & Diaconis, 1981) rule. Dark gray ticks on the abscissa mark the locations of individual data points.

## Logistic regression analysis of image features

We also performed the analysis of image features using logistic regression. Specifically, we used logistic regression to predict the trial by trial responses from the respective distance measure. We then used deviance to quantify how well the respective logistic regression model (and thus the corresponding image feature) explained the observer’s responses. Deviance is a generalization of the sum of squares error to the setting of logistic regression. Under the null-hypothesis that the residuals are simply due to random fluctuations, deviance has a chi-square distribution with degress of freedom equal to the difference between the number of data points and number of parameters in the model.

Figure 9 shows deviances for the different image features. Overall, the results were similar to the analysis based on ROC curves: Radial distances did not predict trial by trial errors very well, while Euclidean distance and Cosine distance in latent space did. There was a significant difference between latent and pixel space for Cosine distance (*t*(6) = −3.86, *p* < 0.05 corrected), but not for Euclidean distance (*t*(6) = −2.46, *p* = 0.048 uncorrected).

**Figure 9:**
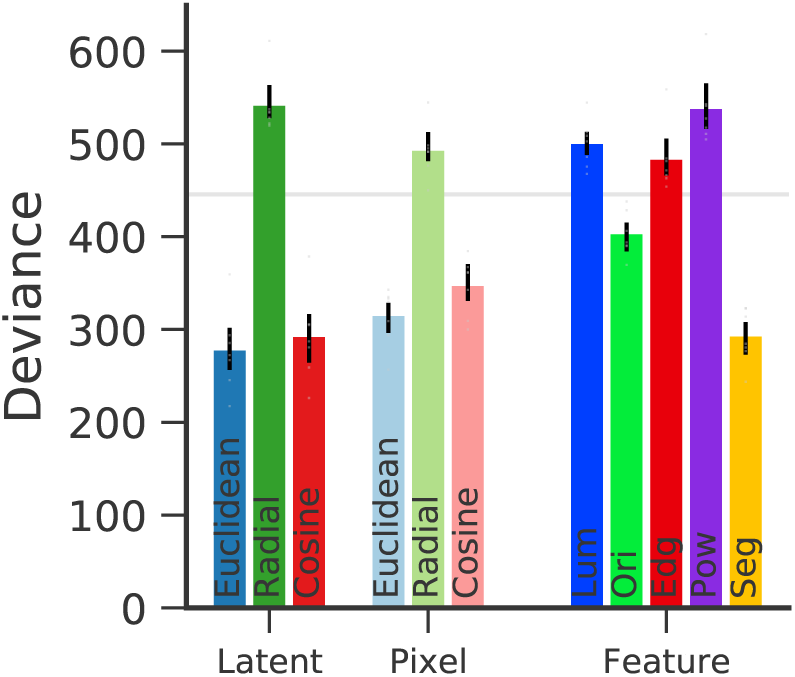
Logistic regression analysis of trial by trial errors. Colored bars indicate mean deviance across observers, error bars are 95% confidence intervals based on bootstrap across observers. Single observer deviances are marked by light gray dots. The horizontal gray line marks the 95-th percentile of the distribution of deviances expected if residuals were due to chance.

Results for different image features were also very similar, with segmentation structure being most predictive. Low level features were generally not much better than chance performance.

